# Machine Learning Assisted Optimization Framework for Designing Transcranial Focused Ultrasound Phased Array Transducer for Deep Brain Neuromodulation in Mice

**DOI:** 10.64898/2026.05.21.727023

**Authors:** Sadman H. Labib, Jingfei Liu

## Abstract

Transcranial focused ultrasound is an emerging noninvasive neuromodulation technique offering high spatial precision and deep penetration. However, in deep brain neuromodulation in mice, the skull base attenuates the signal, distorting the focal region and creating off-target peaks. This study presents a machine-learning–driven simulation framework to optimize a bowl-shaped phased-array transducer design for hypothalamic targeting and compares its performance with that of time-reversal phase conjugation and a single-element baseline. A computed tomography–based mouse head model was used for full-wave acoustic simulations with a fixed bowl geometry (10 mm aperture, 6 mm radius of curvature). Designs were evaluated across various parameters, including operating frequency (0.2–1.5 MHz), active element count (16, 32, 64, 128), and element diameter (300–550 μm). The evaluation employed four metrics: the presence of a −3 dB focal region within the hypothalamic area, axial focal length defined by the −3 dB full-width at half maximum, focal fragmentation measured by the −3 dB blob count, and targeting displacement. Random Forest surrogate models were trained in simulation outputs and paired with the Non-dominated Sorting Genetic Algorithm II to reduce computational costs during multi-objective optimization. The forward-excitation-optimized phased-array design (0.73 MHz, 128 elements, 381 μm element diameter) achieved a focal region at the hypothalamic target with a full width at half maximum of 0.67 mm, a blob count of 1, and a targeting displacement of 0.38 mm when placed 1 mm below the nominal position. Time-reversal phase conjugation further improved confinement and targeting (full width at half maximum: 0.59 mm; displacement: 0.37 mm). Limitations include reliance on a single mouse anatomy, and incorporating additional CT-derived anatomies should enhance generalizability across strains, ages, and sexes.

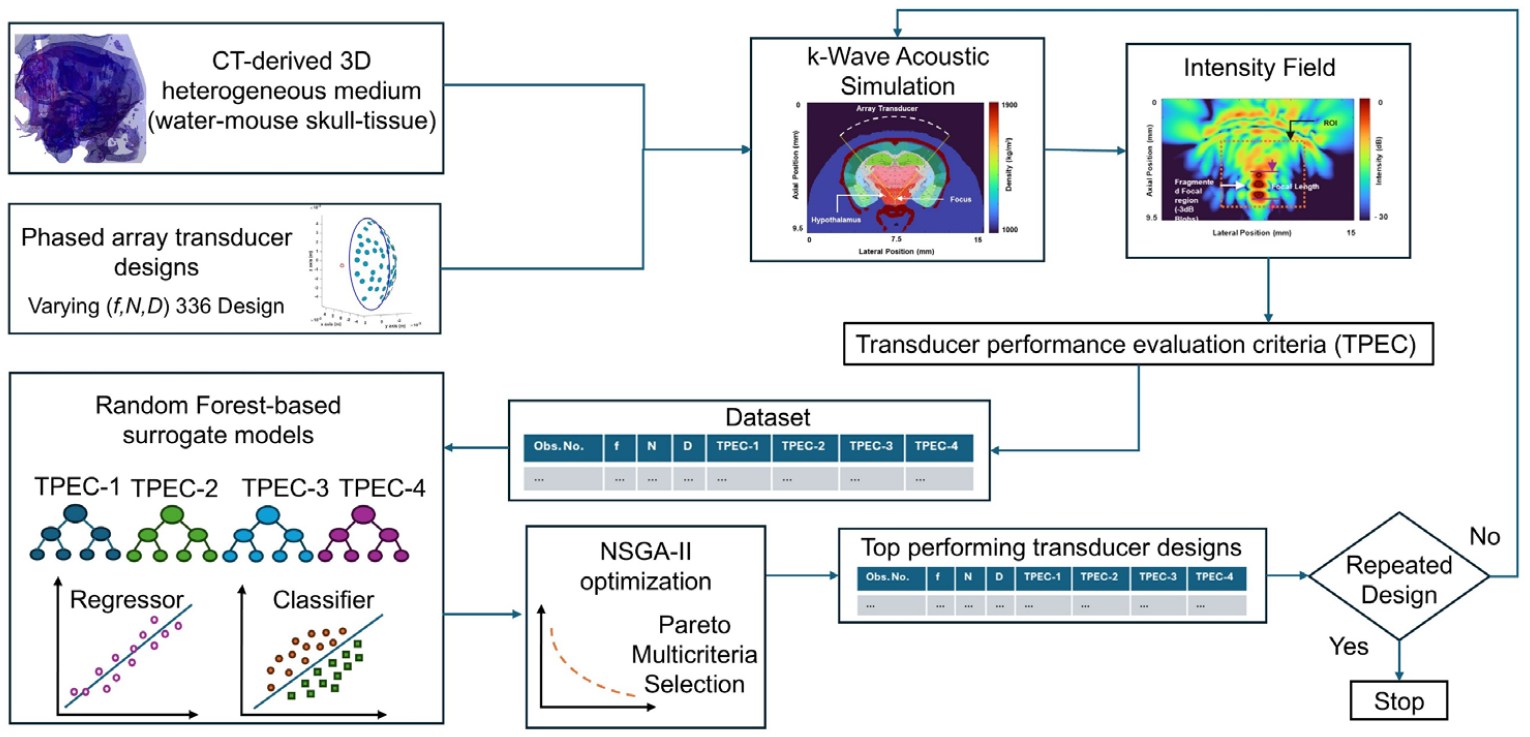

**Highlights:**

- A CT-based acoustic simulation and machine-learning framework was developed to optimize bowl-shaped phased-array transducers for mouse hypothalamic tFUS neuromodulation.
- Random Forest surrogate models coupled with NSGA-II efficiently identified optimized array designs across frequency, element count, and element diameter.
- The optimized phased-array design produced a compact hypothalamic focus with submillimeter targeting displacement, with further confinement achieved using time-reversal phase conjugation.

## I. Introduction

Transcranial focused ultrasound (tFUS), is an emerging noninvasive neuromodulation technology that uses acoustic energy to modulate brain activity with high spatial selectivity and substantial depth penetration [1-3]. Unlike conventional noninvasive brain stimulation modalities such as repetitive transcranial magnetic stimulation, transcranial direct current stimulation, and transcranial alternating current stimulation, which primarily affect superficial cortical regions [4], tFUS can target deep brain circuits and nuclei while operating at low intensities that avoid thermal elevation or mechanical tissue damage [5]. This capability has stimulated rapidly growing interest across neuroscience and medicine, including applications in neurology [6] and psychiatry [7, 8] and in studies of brain function [9, 10] and neurological disorders [11], where the delivery of focused mechanical pressure to neuronal membranes and ion channels drives neuromodulatory effects [12].

For transcranial neuromodulation in rodents, focusing is commonly achieved either with fixed-focus, single-aperture transducers (e.g., curvature-based designs) or with multi-element phased arrays that enable electronic focusing and steering via element-wise phase and amplitude control [13, 14]. Phased arrays can improve spatial control for deep targets and, in some configurations, support multi-focus stimulation strategies [15, 16]. A central technical challenge is the mouse skull: heterogeneous skull structure attenuates and distorts ultrasound, degrading focal accuracy and promoting off-target maxima. Phased arrays can partially mitigate these effects by adaptive phase/amplitude control (aberration correction) to recover focal quality through the skull [17, 18]. However, arrays also introduce practical limitations, including grating and side lobes, increased system complexity, and form-factor constraints (array mass and cabling) that can be particularly restrictive for small-animal behavioral experiments, where maintaining stable coupling and alignment in awake, freely moving subjects is challenging and may necessitate head fixation or anesthesia [19].

Currently, in small-animal t-FUS neuromodulation, most studies have focused on superficial brain regions such as the cortex [20-22], where the acoustic path length through the skull is shorter and skull-induced attenuation and aberration are typically less severe. These shallow targets generally reduce sensitivity to skull heterogeneity and enable more predictable wave propagation and alignment. In contrast, deep-brain targets in mice pose additional technical challenges due to longer propagation distances through heterogeneous skull and brain tissue, increasing cumulative attenuation and aberration. Additionally, for targets near the skull base, the reflected wave front from the internal skull-tissue interface interferes with the converging wavefront, contributing to focal distortion and fragmentation.

A major barrier to advancing preclinical tFUS in freely behaving mice is the lack of a compact transducer design strategy that reliably generates a well-confined, deep focus through the skull while minimizing off-target maxima near the skull or scalp. To address the off-target grating/side-lobe risks that can arise in planar arrays, particularly when electronic steering is required, we adopt a bowl-shaped (geometrically focused) phased-array architecture that concentrates energy toward the intended focal region while retaining element-wise control for transcranial operation. Moreover, phased-array excitation enables tighter, more controllable focusing at moderate frequencies, mitigating the relatively large focal regions typically produced by single-element transducers at low frequencies without incurring the increased attenuation penalties associated with higher-frequency operation.

Here, we develop a CT-derived, anatomically realistic mouse head model and use full-wave simulations to evaluate the performance of bowl-shaped phased arrays across operating frequency (0.2–1.5 MHz), active element number (16, 32, 64, 128), and element diameter (300–550 µm), using a fixed bowl geometry (10 mm aperture, 6 mm radius of curvature) selected from our prior single-element design study [13]. We quantify performance using four metrics: region of interest (ROI) focus presence, axial focal length (−3 dB full length half maximum (FLHM)), focal fragmentation (−3 dB blob count), and targeting or global-peak displacement (to capture off-target maxima relevant to superficial exposure), and train machine-learning surrogate models to enable multi-objective optimization via Non-Sorting Genetic Algorithm (NSGA-II). The resulting forward (FW)-optimized design is benchmarked against time-reversal phase-conjugation focusing as a skull-compensated reference, a single-element baseline, and a placement-sensitivity analysis to quantify robustness to millimeter-scale positioning error.

## II. Methods

When This section describes the simulation, data acquisition, machine-learning-based surrogate modeling, and optimization workflow used to design and evaluate the performance of phased-array t-FUS transducers.

### A. Simulation

In this study, we adopted k-Wave, a widely used open-source MATLAB toolbox for acoustic simulation [23]. We simulated the propagation of ultrasound waves from a bowl-shaped phased array transducer through an inhomogeneous medium (water, bone, and tissue) and ultimately focused them on a deep-brain location. This simulation models the propagation of ultrasound waves through an inhomogeneous medium, accounting for key material properties, including the speed of sound and mass density. The purpose of the simulation is to understand how ultrasound beams are affected by the transducer’s operating frequencies, the number of array elements, and the array element diameters.

#### 1) CT-Based Mouse Head Model

A 3D mouse head model was reconstructed from coronal CT slices obtained from the IMAIOS Vet-Anatomy repository [24]. The CT dataset corresponds to a normal female Swiss mouse acquired using a laboratory microCT system (nanoScan PET/CT, Mediso, Budapest, Hungary) at CERIMED (Aix-Marseille Université, Marseille, France). Acquisition parameters reported by the provider include a tube voltage of 50 kVp and the use of an iodinated contrast agent (Visipaque, 320 mg I/mL; intravenous injection). The stack of 41 coronal slices was reconstructed into a volume and modeled as a 3D simulation medium in k-Wave, comprising 256×169×232 grid points with voxel spacing of (0.058, 0.056, 0.064) mm, corresponding to physical dimensions of (15.0×9.5 ×15.0) mm. The medium is further divided into three regions: water, skull, and tissue, as depicted in Fig. 1, according to their distinct mass densities. Please note that the mouse brain and skin are modeled as the same medium, referred to as soft tissue, because of their negligible differences in acoustic parameters. Assuming each region is homogeneous, the material properties listed in Table I are assigned to the corresponding regions, forming a simplified model of the mouse brain as shown in Fig. 1. In this model, the average skull thickness of the mouse is approximately 0.3 mm [25, 26]. The power law absorption exponent is set to 1.08 [27] as a scalar input, since k-Wave allows only a single value across the entire model.

**TABLE I.** Material Properties Adopted in Simulating the Transcranial Ultrasound Energy Delivery: Speed OF Sound (*c*), Mass Density (*ρ*), Power Law Absorption Prefactor (*α*), power Law Absorption coefficient (*y*)

| Medium | $c$ (m/s) | $\rho$ (kg/m<sup>3</sup>) | $\alpha$ (dB/MHz<sup>γ</sup> cm) | $\gamma$ |
| --- | --- | --- | --- | --- |
| Water [28] | 1500 | 1000 | 0.0022 | 1.08 |
| Skull [29, 30] | 2425 | 1933 | 8.8400 | 1.08 |
| Tissue [30] | 1540 | 1040 | 0.4160 | 1.08 |

**Fig. 1.**
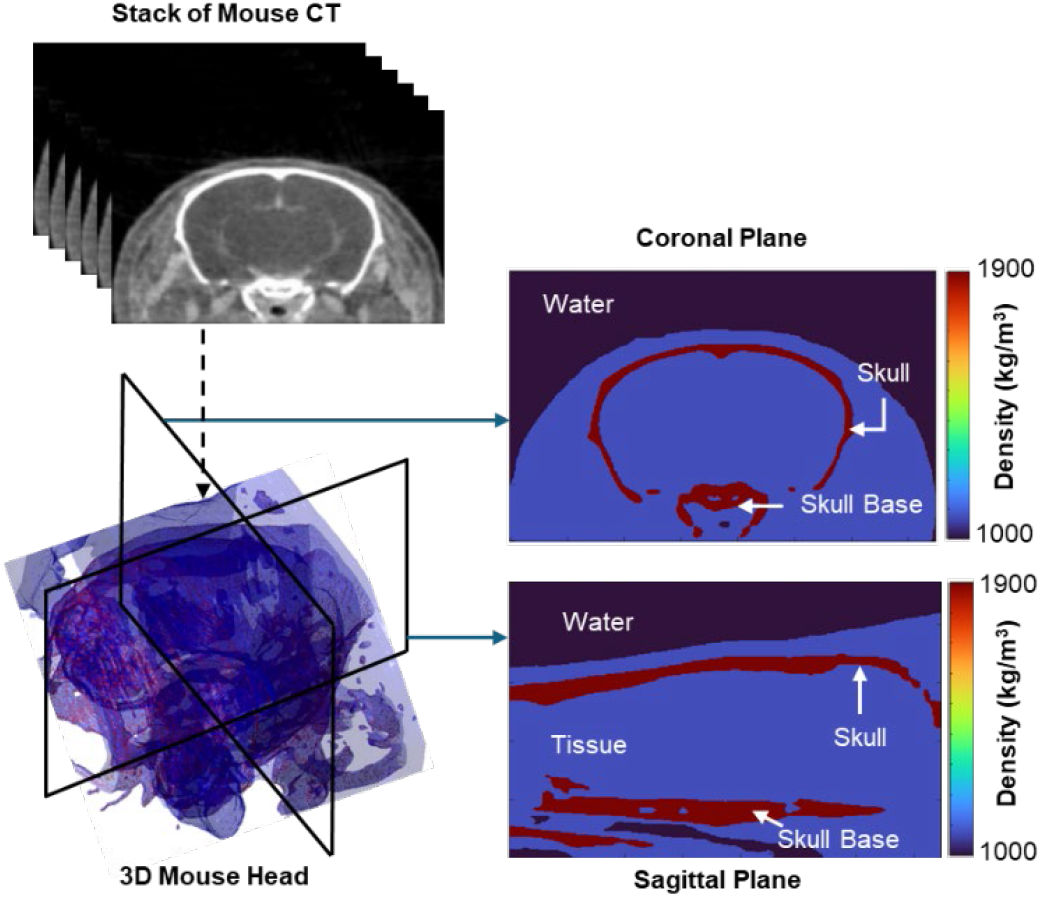
CT-derived mouse head model and segmentation. Coronal CT slices were stacked to reconstruct a 3D mouse head volume and segmented into water (background), tissue, and skull to generate a heterogeneous acoustic model. Representative coronal (top-right) and sagittal (bottom-right) cross-sections highlight the skull and skull base.

**Fig. 2.**
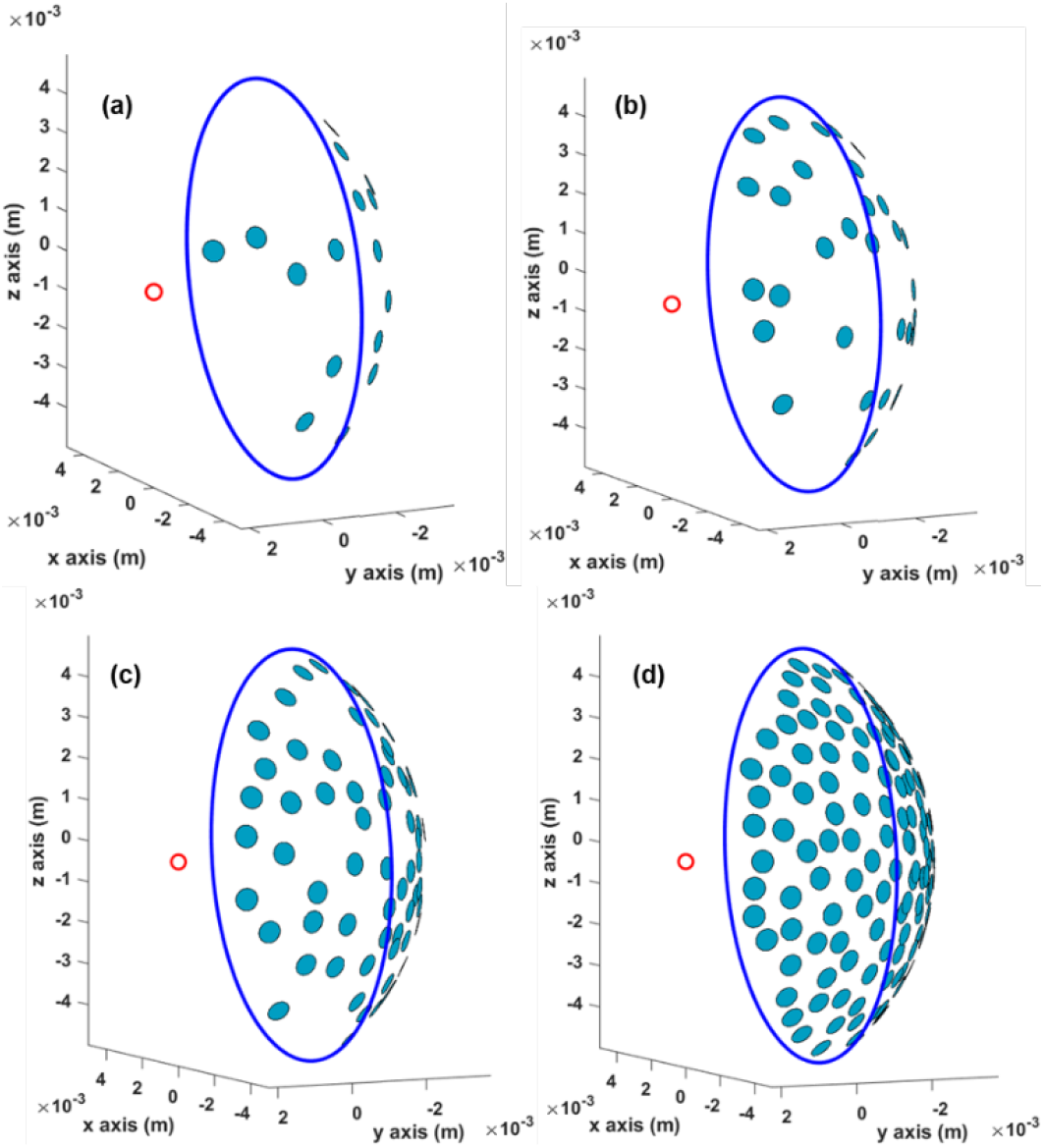
Bowl-shaped phased-array layouts with varying element count. Spherical-cap (bowl) array geometries on a 10 mm aperture and 6 mm radius of curvature showing element center locations for (a) 16 elements, (b) 32 elements, (c) 64 elements, and (d) 128 elements. The blue curve denotes the bowl aperture boundary, and the red marker indicates the geometric focus.

#### 2) Phased array configuration on a bowl-shaped focused transducer

A bowl-shaped phased-array transducer was modeled as a spherical cap with an aperture diameter of 10 mm and radius of curvature (ROC) of 6 mm (Fig. 3). Arrays were designed by varying: (i) the number of active elements, *N* (16, 32, 64, 128) and (ii) the circular element diameter, *D* (300, 350, 400, 450, 500, 550 µm). Each element was represented as a rigid, baffled circular piston (disc) mounted on the spherical cap and oriented toward a prescribed geometric focus (focus position). For all configurations, element centers were restricted to lie on the spherical surface, within the aperture boundary defined by the maximum polar angle (*θ*_max_).

**Fig. 3.**
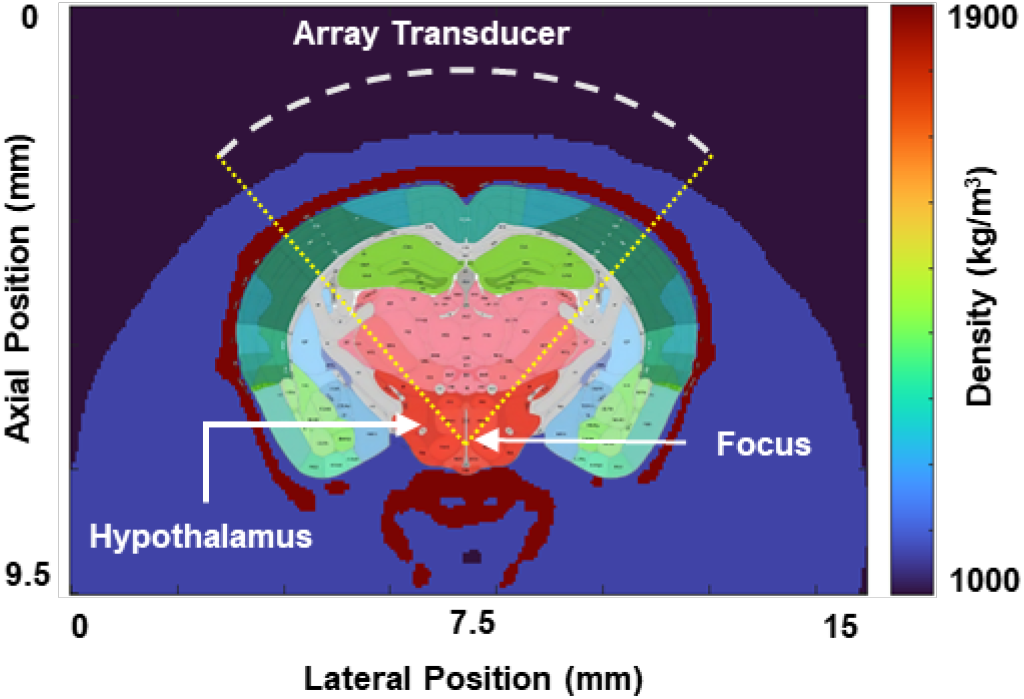
Transducer placement and hypothalamus targeting in the mouse head model.

**Fig. 4.**
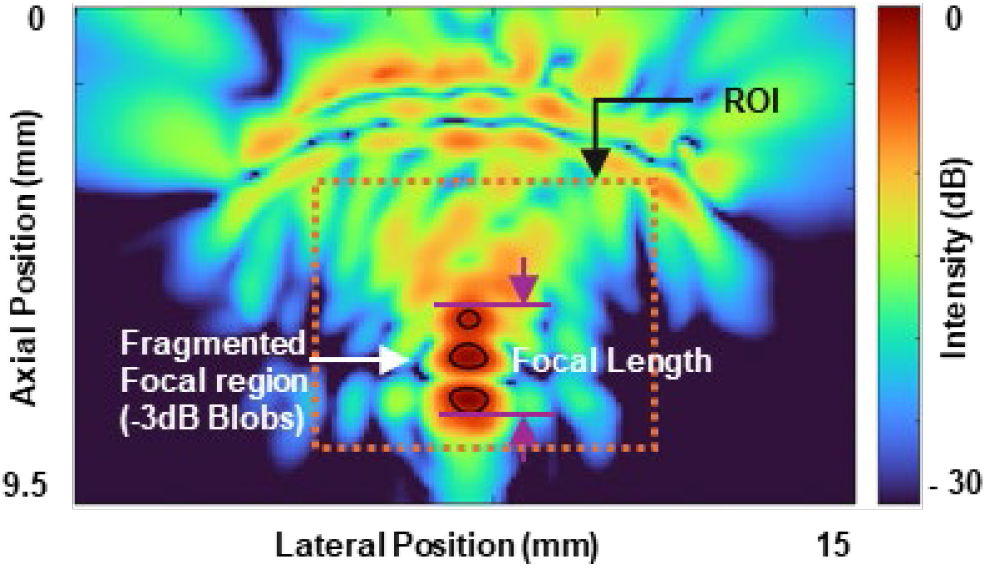
Illustration of the parameters used for the transducer performance evaluation. Showing the focal length (FLHM) above -3 dB, and the fragmented focal region (-3 dB blobs).

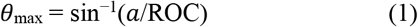

where *a* is the aperture radius.

To generate element layouts without overlapping and with approximately uniform surface coverage, element centers were sampled randomly over the spherical cap using an area-uniform distribution. Specifically, the azimuthal angle was drawn from *ϕ*∼*U*(0,2π), and the polar angle was drawn via

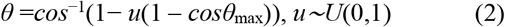

which yields uniform sampling over the cap area. Each sampled direction defines a surface normal 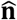 and an element-center position 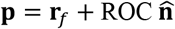, where **r**_*f*_ denotes the focus position. To prevent geometric crowding and enforce manufacturable spacing, a minimum center-to-center distance criterion was imposed:

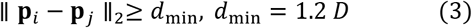

Here *d*_min_ is the minimum center-to-center element distance. Candidate points violating this constraint relative to previously accepted elements were rejected and resampled. Sampling continued until the target element count was reached (or a conservative maximum trial limit was exceeded), ensuring deterministic generation of non-overlapping, well-spaced layouts for each (*N, D*) design. The resulting element positions were instantiated using the ‘`kWaveArray`’ function by adding disc elements at the computed centers with diameter *D* and with each disc assigned an orientation toward the geometric focus. The final array geometry was visually verified by a 3D rendering of the assembled spherical-cap array and the focus point to confirm correct aperture coverage, curvature, and focusing convention.

#### 3) Beamforming Method

Beamforming was implemented using (i) forward continuous wave (FW) simulations to generate the performance dataset for surrogate modeling and optimization, and (ii) time-reversal (TR) beamforming to provide a skull-compensated benchmark for the optimized design. Simulations were performed at operating frequencies (*f*) 0.2 - 1.5 MHz in increments of 0.1 MHz; TR corrections were computed independently at each *f*. Since the aperture and ROC of the array transducer’s bowl is fixed, the transducer position does change during the simulations, varying the (*N*, and *D*. Parameters relevant to k-wave simulations are summarized in Table II.

**TABLE II.** Simulation Parameters for K-Wave Simulations.

| SIMULATION PARAMETERS FOR K-WAVE SIMULATIONS |  |
| --- | --- |
| Simulation Parameters | Value |
| Grid Size ($N_x, N_y, N_z$) | $256 \times 169 \times 232$ |
| Grid Spacing ($d_x, d_y, d_z$) | $0.058 \times 0.056 \times 0.064$ |
| Perfectly Matched Layer (PML) | 10 |
| Min. Points Per Wavelength (PPW) | 18 based on $c_{\text{tissue}}$ |
| Max. Points Per Wavelength (PPW) | 29 based on $c_{\text{skull}}$ |
| Courant–Friedrichs–Lewy number (CFL) | 0.3 |
| Time Resolution (dt) | 6.95 ns |

In FW simulations, all the elements act as the source, andthe k-Wave function ‘`createCWSignals`’ was used to simulate wave signals at 1 kPa pressure. The entire simulation domain was designated as the sensor, and the ‘kspaceFirstOrder3D’ function was used to simulate wave propagation in the time domain. At each time point, the sensor calculated and stored the acoustic pressure and intensity for subsequent analysis.

To achieve deep brain stimulation, the geometric focus (the center of curvature) of the transducer was set to a deep-seated structure located at (7.5 mm, 2.25 mm) in the lateral and axial directions, as shown in Figure 3. This focal arrangement was selected to ensure anatomical relevance in deep-brain murine neuromodulation studies, in which brain regions such as the hypothalamus and brain stem are of interest. The simulated acoustic fields were presented using the root-mean-square pressure (rms pressure, *P*_*RMS*_) and intensity (*I*_*RMS*_). *P*_*RMS*_ was obtained from the time-varying pressure distribution, and *I*_*RMS*_ was calculated using the following relation:

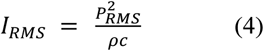

The computed intensity field was first normalized to its maximum value and then converted to a decibel (dB) scale to facilitate visual interpretation and comparative analysis. This approach enabled the identification of high-intensity focal regions and evaluation of off-target energy exposure, including energy deposition within the skull bones and adjacent tissues.

In TR beamforming, a continuous ultrasound wave signal was emitted from a virtual point source located at the geometric focus inside the brain and propagated through the heterogeneous skull–tissue model. During this step, all array elements acted as receivers, and the pressure time series at each element mask point (*i*) was recorded. For each recorded signal, the complex response at the corresponding *f* was extracted using the ‘`extractAmpPhase`’ function, yielding an amplitude *A*_*i*_ and phase *φ*_*i*_ . Time-reversal was implemented via corresponding frequency phase-conjugation, such that 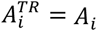 and 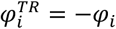 . Then, the phase-conjugation was coupled to the ‘`createCWSignals`’ function and assigned to the same array elements. A second k-Wave simulation was executed using TR-driven wave propagation, and the full-domain *P*_*RMS*_ field was recorded, and *I*_*RMS*_ was subsequently computed, yielding skull-compensated focusing on the intended target by mitigating skull-induced phase aberrations.

### B. Performance Evaluation for Transducer Designs

The goal of this study is to optimize the design of a bowl-shaped phased-array transducer for deep-brain stimulation in mice. By sweeping 14 operating frequencies, 4 element counts, and 6 element diameters, resulting in 336 FW simulated designs, is the initial step of the design optimization process. The mouse deep brain seated hypothalamus was selected as the region of interest (ROI) because it is a representative deep target in the mouse brain and is located approximately 1.5-2.0 mm above the lower skull bone [31]. For each design, we evaluated the simulated intracranial acoustic intensity field and extracted a set of standard focal-quality and targeting metrics to enable consistent, quantitative comparison across the full parameter space. The performance evaluation criteria used in this study are listed below.

1. **Presence of a -3dB focal region at the ROI** Across the 336 simulated designs, a subset of configurations did not produce a contiguous focal region above the -3 dB level within the ROI, indicating that insufficient acoustic energy reached the target due to strong skull-induced attenuation and/or defocusing. We therefore included the existence of a -3 dB region in the ROI as a binary feasibility metric. Specifically, the metric was assigned a value of 1 if at least one connected -3 dB region was present within the ROI (defined relative to the field’s global peak), and 0 otherwise.
2. **Focal Length** Focal length was defined as the axial size of the -3 dB focal region, i.e., the -3 dB full-length at half maximum (FLHM), and reported in millimeters. This metric is a key indicator of axial confinement and potential off-target exposure, because elongated focal regions deposit energy over a larger volume along the propagation direction. As illustrated in Fig. 3, FLHM was computed for each design by identifying the -3 dB region and measuring its extent along the acoustic axis, with the measurement procedure adapted for distorted or fragmented focal shapes. Shorter focal lengths were preferred because they provide tighter axial localization and improved targeting specificity.
3. **Focal Fragmentation** Because the target focus is located near the lower skull bone, a portion of the incident acoustic energy can reflect from the bone-tissue interface and interfere with the incoming field, leading to focal distortion. In several simulations, this interference caused the -3 dB focal region to fragment into two or more spatially separated connected components (blobs). We therefore quantified focal fragmentation by counting the number of connected -3 dB regions within the ROI. A single connected -3 dB region (blob count of 1) was considered desirable, whereas multiple blobs (>1) indicated a distorted/split focus and reduced targeting specificity.
4. **Targeting displacement** Targeting displacement was defined as the Euclidean distance between the prescribed geometric focus (target location) and the location of the global maximum intensity in the simulated field. This metric quantifies steering/focusing accuracy; an ideal design yields a targeting displacement of 0 mm, indicating that peak energy deposition occurs at the intended target.

### C. Machine learning guided Surrogate Modeling

Using the four performance metrics defined above, each of the 336 forward (FW) simulations was post-processed to construct a supervised-learning dataset. The input features comprised the three transducer design variables *f*, N and D, and the outputs comprised the corresponding performance measures: (i) presence of a -3 dB focal region within the ROI (binary), (ii) focal length (-3 dB FLHM, mm), (iii) focal fragmentation based on the -3 dB connected-component count, and (iv) targeting displacement (mm). For classification, the blob metric was binarized to indicate whether a single connected -3 dB region was present in the ROI (1 = single blob, 0 = otherwise).

Surrogate models were implemented in Python using scikit-learn. Two probabilistic Random Forest (RF) classifiers were trained for outputs (i) and (iii) using 1200 trees, with min_samples_leaf = 2 and class_weight = balanced_subsample to mitigate class imbalance. Classifier probabilities were calibrated via isotonic regression with 3-fold cross-validation. For continuous outputs, two RF regressors were trained to target displacement and focal length, using 1500 trees with the default split settings. Because focal length is only meaningful when a focus is present, the focal length regressor was trained only on samples with a valid focal length value.

Model performance was evaluated using a held-out 25% test split (random seed = 42), with stratification by the focus-found label for classification. The focus-found classifier achieved an accuracy of 0.9767, and the single-blob classifier achieved an accuracy of 0.9651. For regression, the targeting-error surrogate achieved the coefficient of determination, *R*^2^=0.854, and the focal-length surrogate achieved *R*^2^ = 0.8733.

### D) Surrogate-Guided Multi-Objective Optimization of Phased-Array Designs

To identify high-performing phased-array designs without additional full-wave simulations, we performed multi-objective optimization using the trained surrogate models as a fast evaluator. The decision vector comprised three design variables: operating frequency *f*, element diameter *D*, and the number of active elements *N*. Frequency and diameter were treated as continuous variables within their admissible ranges. The element count was constrained to the discrete set [16, 32, 64, 128] by optimizing an index variable and decoding it to the nearest allowed value during evaluation.

Optimization was carried out with the NSGA-II evolutionary algorithm implemented in ‘pymoo’ using a population size of 100 and 60 generations (random seed = 42). For each candidate design **x** = [*f, N, D*], the surrogate models returned: (i) *P*_*focus*_, the predicted probability that a -3 dB focal region exists within the ROI, (ii) *P*_*single*_, the predicted probability that the ROI contains a single connected -3 dB region (i.e., no focal splitting), (iii) the predicted focal length (FLHM), and (iv) the predicted targeting displacement. A single feasibility constraint was imposed to exclude designs unlikely to form a focus at the ROI:

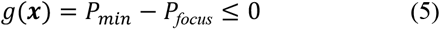

where *P*_min_ = 0.70 was used as a conservative threshold. Among feasible designs, NSGA-II minimized three objectives:

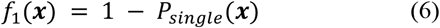

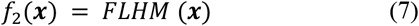

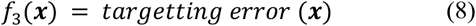

This formulation promotes designs that are likely to generate a single, compact focus with minimal displacement in targeting, while ensuring a high probability of focus formation within the ROI. The resulting feasible, non-dominated (Pareto-optimal) solutions were extracted from the final population, deduplicated, and exported for reporting and subsequent full-wave validation.

The top-performing designs identified by NSGA-II were subsequently re-simulated using k-Wave to obtain performance metrics. These newly simulated samples were appended to the existing dataset, and the surrogate modeling and optimization steps (Sections C and D) were repeated for two additional rounds. This iterative surrogate–simulation refinement continued until the optimization converged, defined here as the recurrence of the same top-ranked design(s) across successive iterations (Fig. 5).

**Fig. 5.**
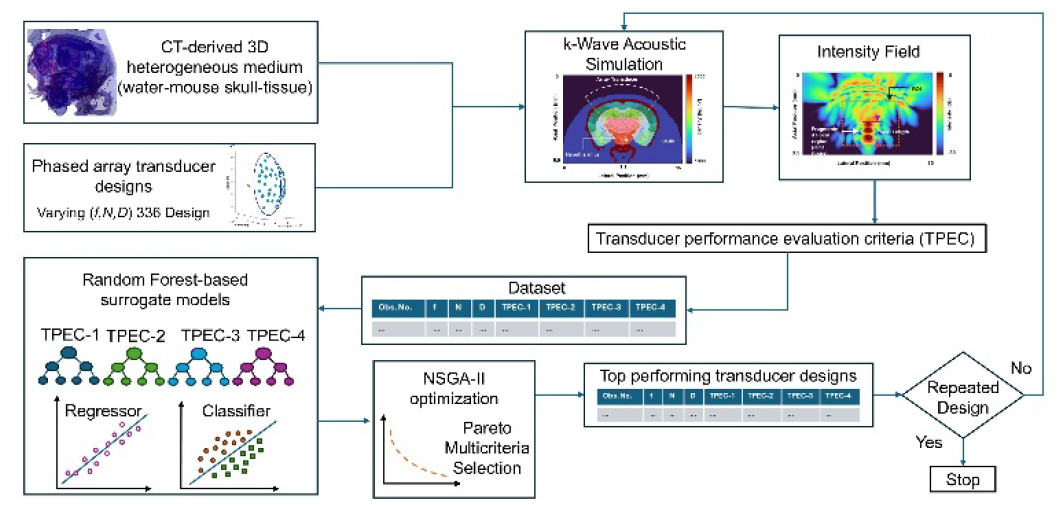
End-to-end workflow for CT-guided phased-array transducer optimization. A CT-derived mouse head model and a bowl-shaped phased-array geometry are used to perform full-wave k-Wave simulations over a sweep of design parameters (frequency, number of elements, and element diameter). Simulated fields are reduced to transducer performance evaluation criteria (TPECs), including ROI focus presence, axial focal length (FLHM), focal fragmentation (−3 dB blob count), and targeting displacement, and compiled into a structured dataset. Random Forest surrogate models are trained on this dataset and coupled with NSGA-II multi-objective optimization to identify top candidate designs. The highest-performing designs are re-simulated and appended to the dataset, and the surrogate optimization loop is repeated until the top optimized design converges (repeat selection), after which the final design is reported.

## III. Results

### A. Surrogate Model Validation and Feature Sensitivity

Surrogate performance was evaluated on a held-out 25% test split (random seed = 42), with stratification by the focus-found label for classification. The focus feasibility classifier (Presence of a -3dB focal region at the ROI) achieved an accuracy of 0.9667, an F1-score of 0.96, and a Receiver Operating Characteristic–Area Under the Curve (ROC–AUC) of 0.9934. The single-blob classifier (Focal fragmentation) achieved an accuracy of 0.9556 and ROC–AUC of 0.989, with a lower F1-score of 0.7143, consistent with the limited number of single-blob outcomes in the dataset. For regression, the targeting or global-peak displacement surrogate achieved a mean absolute error (MAE) of 0.456 mm, a root mean squared error (RMSE) of 1.3491 mm, and a coefficient of determination (*R*^2^) of 0.7648. The axial focal length surrogate (focal length), trained only on cases where a focus existed at the ROI (focus_found = 1), achieved MAE = 0.4409 mm, RMSE = 1.0322 mm, and *R*^2^ = 0.5712. Permutation feature importance (Fig. 6) indicated that element number was the dominant contributor to predicting focus feasibility and global-peak displacement, whereas operating frequency most strongly influenced axial focal length and the probability of obtaining a single connected −3 dB focal region; element diameter showed a smaller secondary effect across tasks.

**Fig. 6.**
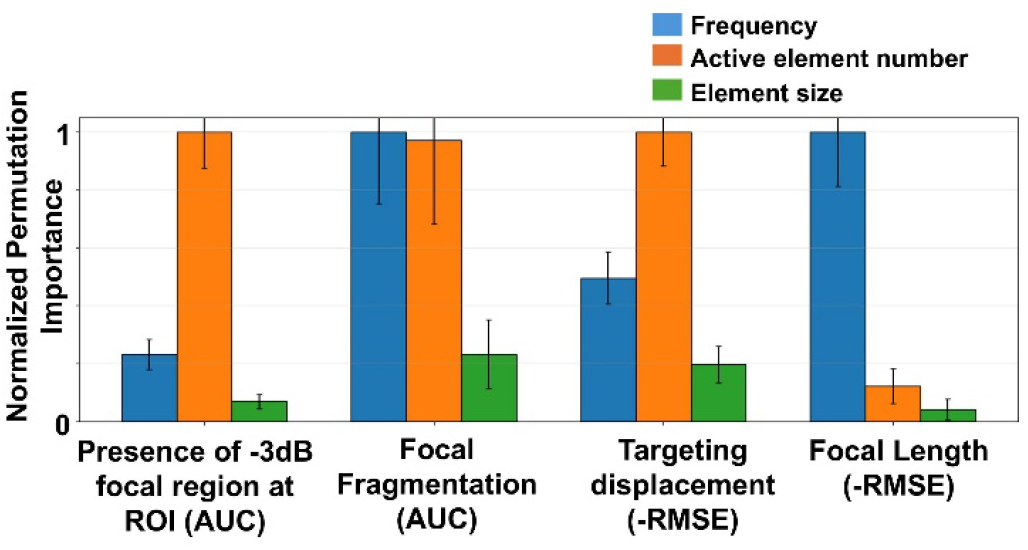
Permutation feature importance for surrogate models. Normalized permutation importance (mean ± standard deviation across permutations) of the three design variables: *f, N*, and *D* for predicting (i) focus presence in the ROI and (ii) single-blob occurrence (reported using ROC–AUC), and for regressing (iii) global-peak displacement and (iv) axial focal length (RMSE). Higher values indicate greater sensitivity of model performance to the corresponding variable.

### B. Optimized Design Transducer

Surrogate-guided multi-objective optimization converged to a single preferred phased-array configuration for detailed evaluation: *f* = 0.73 MHz, *N* = 128, and *D* = 381 μm (aperture diameter 10 mm, ROC 6 mm). FW simulation confirmed the presence of a -3 dB focal region within the hypothalamic ROI, Fig. 7 (a) and (b). The FW intensity field exhibited an axial focal length of FLHM 1.96 mm, two connected -3 dB regions within the ROI (blob count of 2), and a global-peak displacement of 0.37 mm relative to the intended target. The pressure at the focus was 0.63 MPa, and the peak pressure in the field was 0.82 MPa. Importantly, the global maximum occurred adjacent to the target region rather than at the skull or scalp boundary, indicating the absence of a superficial pressure hot spot for this configuration under the modeled conditions. For reference, because only *P*_*rms*_ was stored, the Mechanical Index (MI) was estimated for a near-sinusoidal steady-state waveform using 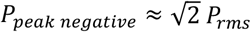 and 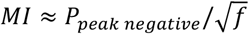. Under this assumption, at 0.73 MHz, the estimated MI was about 1.04 at the target and 1.36 at the global peak.

**Fig. 7.**
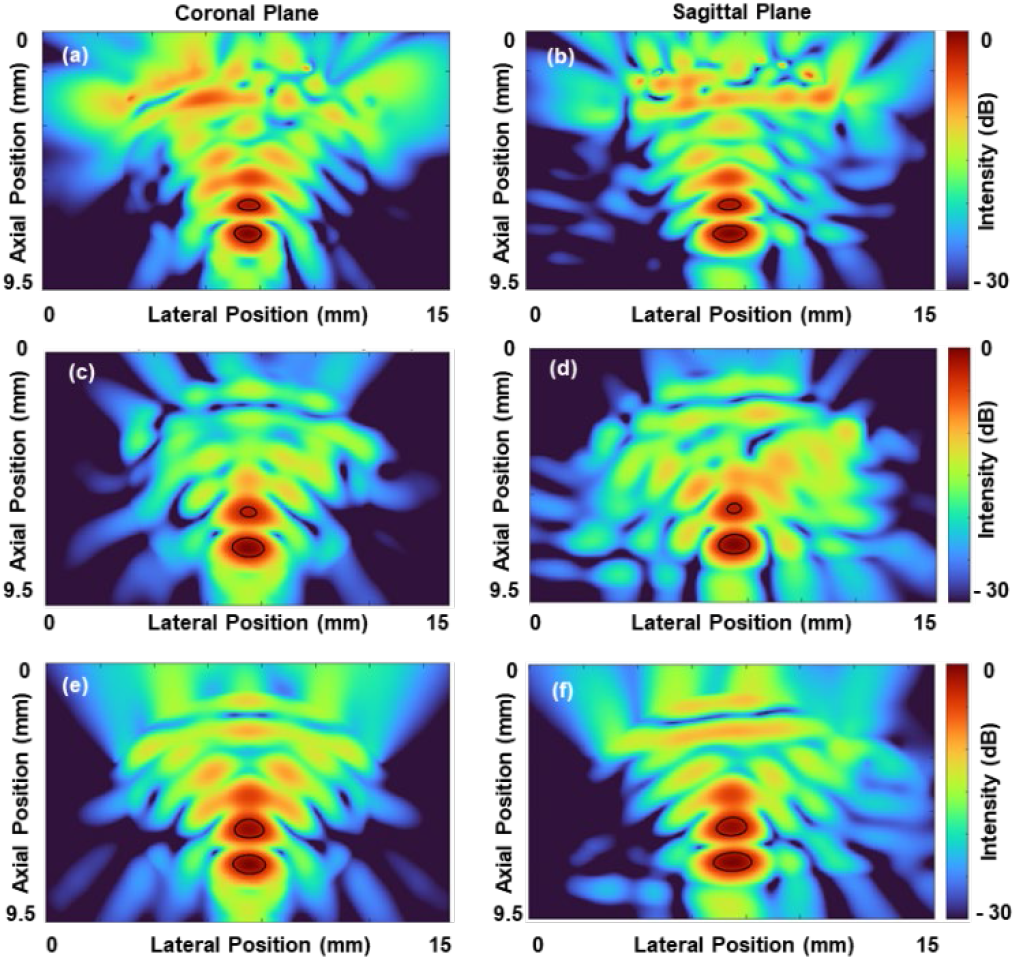
Illustration of the acoustic intensity field with -3dB contour overlaid (black). Optimized phased-array transducer with FW beamforming illustrated in (a) and (b), with TR beamforming in (c) and (d). Single element transducer with same aperture and ROC in (e) and (f).

### C. Comparison with Time Reversal Beamforming

Time-reversal (TR) beamforming (single-frequency phase conjugation at *f* = 0.73 MHz) was used as a skull-compensated benchmark for the selected design. TR preserved focus formation within the ROI and improved both targeting and axial confinement. Specifically, TR reduced the global-peak displacement from 0.37 mm (FW) to 0.30 mm (TR) and FLHM from 1.96 mm (FW) to 1.85 mm (TR), while the -3 dB region remained split into two blobs (blob count: 2). These results, illustrated in Fig. 7 (c) and (d), indicate that, for this geometry and target location, TR primarily improves targeting accuracy, whereas residual focal fragmentation likely reflects local interference effects near the lower skull boundary in addition to phase aberration.

### D. Comparison with a Single Element Focused Transducer

- The optimized phased-array design was benchmarked against a single-element bowl-shaped transducer under the same head model and target definition. The single-element transducer produced a -3 dB focal region at the ROI with FLHM = 2.04 mm, blob count of 2, and targeting displacement of 0.36 mm, Fig. 7 (e) and (f). Compared with the single-element baseline, the optimized phased-array achieved a modest reduction in axial focal length (1.96 mm vs 2.04 mm) under FW excitation and further improved axial confinement under TR (1.85 mm), while maintaining the same focal fragmentation class (two blobs). The global-peak displacement was comparable between the phased-array FW and single-element cases (0.37 mm vs 0.36 mm), and improved under TR (0.30 mm), supporting the benefit of phase compensation for mitigating skull-induced mistargeting.

### E. Best performing Transducer Design from Time-Reversal Simulation

Because time-reversal (TR) beamforming compensates for skull-induced phase aberrations, the transducer configuration that performs best under TR is not necessarily the same design identified by optimization based on forward (FW) excitation. In our FW dataset, cases exhibiting a single connected −3 dB focal region within the ROI were relatively rare, limiting the optimizer’s ability to consistently identify configurations that achieve minimal focal fragmentation under open-loop excitation. Therefore, to quantify the achievable focusing performance under ideal phase correction, we separately screened candidate configurations using TR beamforming and ranked them directly using the TR-derived performance metrics.

For TR evaluation, each design was processed using a two-step workflow: (i) a forward run with a virtual point source at the intended target to obtain the element-wise complex response, followed by (ii) a second run in which phase-conjugated continuous-wave (CW) sources were applied at the same operating frequency to reconstruct a skull-compensated focus. The resulting TR fields were assessed using the same metrics as in the FW analysis: presence of a -3 dB focal region within the ROI, axial focal length (FLHM), focal fragmentation (-3 dB blob count), and global peak displacement.

Among the TR-evaluated designs, the best overall TR performance was obtained for *f* = 1 MHz, *N* = 128, and *DD* = 550 μm. This configuration produced a clear -3 dB focal region within the ROI with a compact axial extent (FLHM = 0.59 mm) and a single connected -3 dB region within the ROI (blob count of 1). The targeting displacement for this TR-focused field was 0.38 mm, indicating accurate skull-compensated focusing relative to the intended target. Collectively, these results (Fig. 8) show that the optimal parameter set depends on the beamforming strategy: FW optimization identifies a practical open-loop design, whereas the TR-selected configuration provides an upper-bound benchmark for achievable focal quality when skull-induced phase distortions are ideally corrected.

**Fig. 8.**
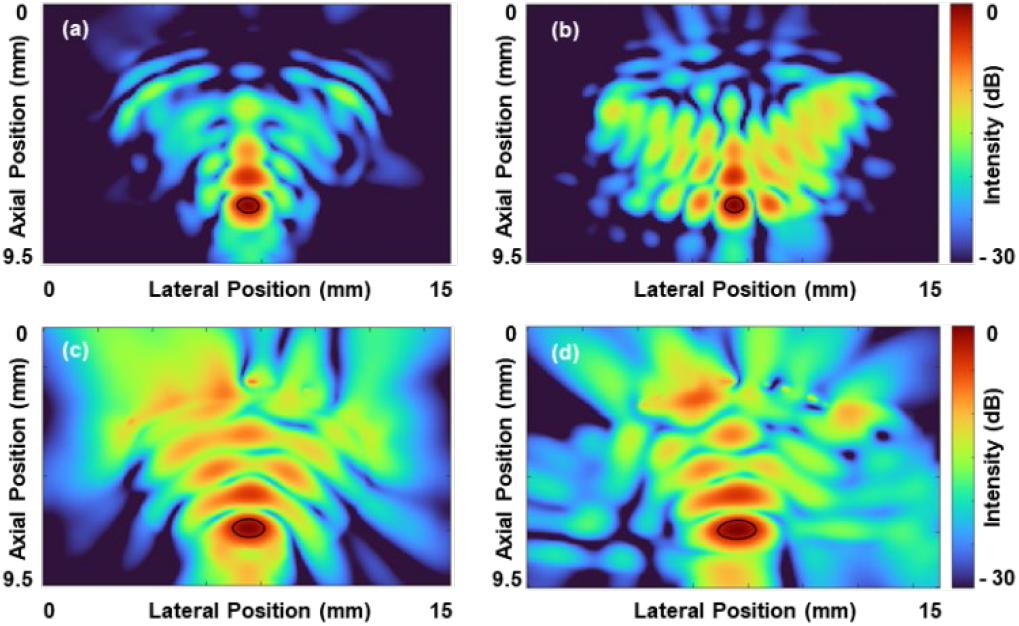
Single-blob focusing under skull-compensated TR beamforming and placement-adjusted FW operation. Intensity field maps (with the -3 dB contour overlaid (black) are shown for the best-performing phased-array configuration under time-reversal (TR) phase-conjugation beamforming in the (a) coronal and (b) sagittal planes. Panels (c) and (d) show the FW-optimized phased-array design after a 1 mm downward translation in the (c) coronal and (d) sagittal planes.

### F. Sensitivity Analysis of Optimized Transducer

To assess robustness to placement variability, the FW-optimized transducer was translated by ±1 mm along each Cartesian direction (up, down, left, right, forward, and backward) relative to the nominal placement, and the four-performance metrics were recomputed, Table III. For all six perturbed positions, a −3 dB focal region remained present within the ROI, indicating that the design is feasible under millimeter-scale placement offsets. Despite preserved feasibility, focal quality and targeting were sensitive to placement direction. The FLHM varied substantially, ranging from 0.67 mm (1 mm down) to 3.15 mm (1 mm up). Similarly, focal fragmentation increased under some offsets, with blob count ranging from 1 (down) to 3 (up). The targeting displacement also increased relative to the nominal case, spanning 0.652–2.1 mm across offsets; lateral shifts (left/right) produced the largest targeting displacement (2.1 mm), while the 1 mm down shift yielded the smallest error (0.652 mm). Forward/backward shifts produced intermediate degradation (errors of 2.0 mm and 1.94 mm, respectively). Overall, these results show that the FW-optimized design is robust in focus formation but exhibits direction-dependent sensitivity in confinement, fragmentation, and targeting accuracy under millimeter-scale placement perturbations.

**TABLE III.** Performance Evaluation Metrics of the Transducer at Different Translated Positions.

| Translation Position of the bowl | Focus at ROI (-3 dB Present) | Focal Length (mm) | Focal Fragmentation (Count) | Targeting Displacement (mm) |
| --- | --- | --- | --- | --- |
| Nominal | 1 | 1.96 | 2 | 0.37 |
| Up | 1 | 3.15 | 3 | 1.30 |
| Down | 1 | 0.67 | 1 | 0.65 |
| Left | 1 | 2.20 | 2 | 2.10 |
| Right | 1 | 2.08 | 2 | 2.10 |
| Forward | 1 | 1.86 | 2 | 2.00 |
| Backward | 1 | 1.91 | 2 | 1.94 |

## IV. Discussion

### A. Effects of Frequency, Element Number, and Element Diameter on Focusing Performance

The present study examined how *f, N*, and *D* govern intracranial focusing through the mouse skull. Across the evaluated parameter space, performance trends were primarily driven by *f* and *N*, with *D* exerting a secondary influence. Frequency controls the trade-off between focal confinement and transcranial transmission: increasing *f* can tighten the focal region in homogeneous media, but in the transcranial setting, it also increases attenuation and phase distortion, thereby reducing delivered energy at the target and promoting off-target maxima. Increasing *N* improves aperture sampling and phase controllability, thereby increasing the likelihood of forming a focus at the ROI and improving targeting accuracy. Changes in *D* mainly affect packing density and spatial sampling on the spherical cap, influencing the interference structure and sidelobes; within the tested range, its contribution was smaller than those of the other two factors. These interpretations are consistent with the surrogate-model feature sensitivity (Fig. 6), which indicated that *N* contributed most strongly to focus feasibility and global-peak displacement, whereas *f* contributed most strongly to focal extent and the probability of obtaining a single connected −3dB region.

### B. Physical Basis for Focal Fragmentation Near the Skull Base

A central challenge in mouse deep-brain targeting is that the desired ROI may lie near strong acoustic-impedance interfaces, such as the lower skull boundary. In this regime, partial reflections generate a superposition between the incident and reflected fields, creating interference patterns that can split the −3dB region into multiple connected components (focal fragmentation). This effect is amplified when the target is within a few wavelengths of the boundary, where constructive interference can occur at multiple nearby locations. In addition, the heterogeneous skull introduces spatially varying transmission and phase delay, further promoting multi-lobe structures rather than a single compact focus. The frequent appearance of fragmented −3dB regions in the FW simulations is, therefore, consistent with the combined effects of (i) boundary-reflection-driven interference near the skull base and (ii) aberration and attenuation introduced by the skull.

### C. Improved performance of Time Reversal Beamforming Compared to FW Beamforming

TR beamforming improves focusing by compensating skull-induced phase aberrations using phase-conjugated element excitations derived from the medium-specific response. This correction typically improves targeting and focal stability compared with open-loop FW excitation, particularly in heterogeneous transcranial propagation. However, TR does not eliminate all degradation mechanisms: fragmentation can persist when the dominant limitation is interference from strong reflectors near the target, rather than phase distortion across the skull alone. Importantly, the design that performs best under TR need not match the FW-optimized design, as TR changes the excitation regime by imposing skull-compensated phases. This distinction is particularly relevant here because single-blob outcomes were relatively scarce in the FW dataset, limiting the optimizer’s ability to reliably converge toward fragmentation-minimizing solutions in open-loop conditions.

### D. Comparison of FW-optimized Design and TR-best benchmark and practical placement guidance

To provide a deployable open-loop design while also benchmarking skull-compensated performance, we report two complementary best-performing configurations. The FW-optimized transducer design selected by machine-learning-assisted optimization was 128 elements, with 381 µm diameter driving at 0.73 MHz. At the nominal placement, this design produced a −3dB focal region within the ROI with FLHM 1.96 mm, two connected −3dB regions, and a global-peak displacement of 0.37 mm. The placement-robust analysis further showed that focus feasibility was preserved under all ±1 mm translation, but focal quality was direction dependent (Fig. 8). Notably, a 1 mm downward translation improved focal compactness and fragmentation, yielding a single connected −3dB region with FLHM 0.67 mm. This improvement was accompanied by an increased global peak displacement (0.652 mm); however, the peak region was located in the hypothalamus. Conversely, upward and lateral translations degraded focal confinement and/or increased fragmentation (e.g., FLHM up to 3.15 mm with three blobs in the 1 mm up case), illustrating that millimeter-scale placement differences can materially alter the focal structure even when a focus remains present within the ROI.

In contrast, the TR-best configuration is reported as a skull-compensated benchmark, representing an upper bound on achievable focal quality under ideal phase correction. Under TR beamforming, 128 elements, with 550 µm diameter driving at 1 MHz delivers a clear −3dB focal region in the ROI with FLHM 0.59 mm, single-blob behavior, and a global-peak displacement of 0.38 mm. Reporting both outcomes clarifies that “best” depends on whether the operating mode is (i) practical open-loop FW excitation, where placement can be used to mitigate fragmentation, or (ii) skull-compensated focusing, where TR provides a useful reference benchmark.

### E. Surrogate-model Performance and Implications for Optimization Outcomes

The surrogate models achieved strong discrimination for focus feasibility and good prediction accuracy for global-peak displacement, supporting their use for efficient multi-objective screening. The focus feasibility classifier achieved an accuracy of 0.9667 and ROC–AUC of 0.9934. The single-blob classifier achieved an accuracy of 0.9556 and ROC–AUC of 0.989, but a lower F1-score (0.7143), consistent with class imbalance due to the scarcity of single-blob outcomes in FW. For regression, the target-error surrogate achieved *R*^2^ = 0.7648, whereas the axial-length surrogate achieved a lower *R*^2^ = 0.5712, suggesting that focal extent is more difficult to predict from limited data when fragmentation and boundary effects introduce additional variability. These results motivate two practical conclusions: the surrogate-guided optimizer can reliably identify feasible designs with improved targeting metrics, but objectives associated with rare outcomes (e.g., single-blob FW focusing) may require iterative dataset enrichment around promising designs to improve predictive fidelity.

### F. Limitation and Future Work

This study has several limitations that motivate future investigation. First, the acoustic model was derived from a single mouse CT dataset and therefore does not capture inter-animal variability in skull thickness, curvature, and heterogeneity; validating across multiple animals is necessary to establish robustness. Second, although the parameter sweep spanned a wide range of *f, N*, and *D*, the FW dataset contained relatively few single-blob outcomes, limiting data-driven optimization for fragmentation-related objectives under open-loop excitation. Third, TR results represent a best-case benchmark requiring element-wise phase control and a subject-specific phase-planning workflow. Future work should therefore (i) repeat the workflow across multiple anatomical models, (ii) iteratively enrich the dataset in regions yielding near-single-blob focusing to improve surrogate reliability for fragmentation outcomes, and (iii) validate the predicted trends experimentally. For longer-term translation, the same pipeline can be extended to clinically relevant skull geometries and design constraints, where lower frequencies and larger apertures with higher element counts are expected to be advantageous for penetration and aberration correction.

## V. Conclusion

This study presented a machine-learning–guided design workflow for mouse transcranial focused ultrasound (tFUS) that combines a CT-derived anatomical head model, full-wave acoustic simulations, and multi-objective optimization to identify compact bowl-shaped phased-array configurations for deep-brain targeting. A FW optimized design: *f* = 0.73 MHz, *N* =128, D= 381µm ; 10 mm aperture, 6 mm radius of curvature) consistently produced a −3 dB focal region at the hypothalamic ROI and achieved nominal performance of FLHM = 1.96 mm, blob count = 2, and global-peak displacement = 0.37 mm, with a target RMS pressure of 0.63 MPa and a global maximum RMS pressure of 0.82 MPa localized near the target rather than at the skull/scalp. A placement-sensitivity analysis showed that focus feasibility was preserved for all ±1 mm translations, whereas focal quality was direction-dependent; notably, a 1 mm downward shift yielded a single-blob focus with FLHM = 0.67 mm.

To benchmark skull-compensated performance, time-reversal (TR) phase-conjugation beamforming was evaluated separately and produced a distinct best-performing configuration *f* = 1 MHz, *N* =128, *D* = 550µm achieving single-blob focusing with FLHM = 0.59 mm and global-peak displacement = 0.38 mm, indicating that the optimal parameter set depends on whether excitation is open-loop or skull-compensated. Surrogate models achieved high predictive performance for focus feasibility (accuracy 0.9667; ROC–AUC 0.9934) and strong performance for global-peak displacement (*R*^2^ = 0.7648), enabling efficient search of the design space and informing sensitivity to the governing design variables.

Overall, the results demonstrate that (i) surrogate-based optimization can efficiently identify feasible mouse tFUS array designs, (ii) skull-induced interference near the skull base is a key driver of focal fragmentation under FW excitation, and (iii) modest placement adjustments can materially improve focal integrity in practical preclinical setups. Future work will extend validation across multiple animals to quantify anatomical variability, enrich the training set to better capture rare single-blob FW outcomes, and incorporate experimental validation and exposure-specific safety analyses to support translation toward robust preclinical and, ultimately, human-relevant systems.

